# Girdin controls the pace of 3D tracheal cell intercalation by coupling adherens junctions to the actin cytoskeleton in *Drosophila*

**DOI:** 10.64898/2025.12.01.691508

**Authors:** Sandra Carvalho, Patrick Laprise, Antoine Guichet, Véronique Brodu

**Author notes:** Co-corresponding authors, **Contact** Correspondence and requests for materials should be addressed to V. B. or A. G., Véronique Brodu, Antoine Guichet, Université Paris Cité, CNRS, Institut Jacques Monod UMR 7592, 15 rue Hélène Brion, 75205 PARIS CEDEX 13, France.

## Abstract

Morphogenesis is orchestrated through coordinated cell movements, including cell intercalation, which drives extensive changes in cell shape and position. This process requires precise regulation of interactions between Adherens Junctions (AJs) and the cortical actin network to generate the necessary mechanical forces. Although the Myosin molecular motor II plays a central role in generating morphogenetic forces, it is dispensable for processes such as dorsal branch (DB) morphogenesis in the *Drosophila* tracheal system, where coordinated cell migration and three-dimensional intercalation rely on alternative, yet poorly characterised mechanisms of force generation. Here, we show that Vinculin and Ajuba LIM protein, two key regulators of mechano-transduction and cell adhesion, do not localise to AJs of DBs, while the cytoskeletal adaptor Girdin does. We demonstrate that Girdin’s function relies on AJ components α-Catenin and E-Cadherin and is specifically required in tracheal cells to ensure the proper pace of cell intercalation. In addition, Girdin contributes to the actin network enrichment at AJs. Its role as an integral AJ component is further elucidated through the development of a novel genetic tool enabling *in vivo* cell-specific actin depolymerisation.

**Summary Statement:** *Drosophila* Girdin contributes to the pace of tracheal cell intercalation by linking adherens junctions to actin. A genetic tool for actin depolymerisation reveals Girdin as a key adherens junction component.

## Introduction

Morphogenetic events are largely driven by coordinated cellular movements that sculpt tissues and the developing organism. These movements are associated with changes in cell shape and position, driven by the cytoskeleton and associated proteins (Friedl & Gilmour, 2009). Intercellular signals coordinate the activities of individual cells to achieve orchestrated tissue movement. However, how cells are coordinated to move remains a significant gap in our understanding of morphogenesis (Friedl & Mayor, 2017; Mishra et al., 2019). Investigating morphogenesis in whole organisms is essential, as it allows for consideration of the contributions of surrounding tissues.

During tissue morphogenesis and repair, AJs are required to both maintain tissue architecture and enable cell movements. The AJ complex, of which cadherin is an essential component, links neighbouring cells. In *Drosophila melanogaster*, E-Cadherin (E-Cad, encoded by *shotgun* (*shg*)) is the only cadherin expressed in embryonic epithelial cells (Tepass et al., 1996) and its highly conserved cytoplasmic tail interacts with α-Catenin and β-catenin to link them to the actin cytoskeleton. This relationship is mutually beneficial: actin supports AJ formation and stability, and AJs contribute to actin network organisation (Campàs et al., 2024).

*In vivo*, mechanical forces on AJs reinforce both homophilic E-Cad adhesion and α-Catenin–F-actin bonds through load-dependent conformational changes that increase the affinity and/or the stability of interacting proteins (Pinheiro & Bellaïche, 2018). E-Cad responds through extracellular domain conformational changes, while α-Catenin catch bonds require both β-catenin and F-actin binding (Pinheiro & Bellaïche, 2018). Under tension, open conformation α-Catenin stabilises F-actin binding and recruits additional actin-binding proteins such as Vinculin (under high tension; le Duc et al., 2010) and Ajuba LIM protein (jub; under lower tension; Sheppard et al., 2023). Vinculin prolongs α-Catenin conformation change and thus tension-dependent AJ reinforcement (Yao et al., 2014). Interestingly, vinculin knockout is deleterious in some organisms, such as worms (Barstead & Waterston, 1991) and mice (Xu et al., 1998). In contrast, zygotic vinculin function has been reported as dispensable in zebrafish (Han et al., 2017), and vinculin is nonessential in *Drosophila* (Alatortsev et al., 1997; Maartens et al., 2016), unlike the major epithelial disruptions observed upon loss of any single AJ component (Sheppard et al., 2023). This difference could be a result of functional redundancy among α-Catenin’s interactors (Pinheiro & Bellaïche, 2018).

The mammalian cytoskeletal adaptor protein CCDC88A, also known as Girdin (Enomoto et al., 2006), promotes anchoring of the E-Cadherin–β-catenin complex to the actin cytoskeleton by interacting directly with actin microfilaments *via* its C-terminal domain 2 (Enomoto et al., 2005). The gene is conserved in *Drosophila*, which expresses a single protein ortholog. Girdin depletion strongly enhances adhesion defects associated with reduced E-Cad expression (Houssin et al., 2015). Moreover, the fraction of E-Cad that co-immunoprecipitates with the actin pool decreases in the absence of Girdin, thereby identifying it as a positive regulator of AJ function (Houssin et al., 2015). In addition, the complete loss of Girdin leads to multiple defects in epithelial tissue morphogenesis and collective cell migration, resulting in dorsal closure defects (Houssin et al., 2015).

The contributions of the AJ-associated cortical actin network have been extensively characterised during two-dimensional (2D) cell intercalation (Alatortsev et al., 1997; Maartens et al., 2016; Pinheiro & Bellaïche, 2018). In 2D, the molecular motor Myosin II (Myo II) associates with the subcortical actin cytoskeleton to generate localised actin contraction, bringing distant cell membranes together and promoting intercalation. Therefore, 2D cell intercalation arises from internal cell forces generated by actomyosin contractile cables (Paré & Zallen, 2020). However, this mechanism accounts for only a small subset of intercalation events in the 3D tissues of developing organisms, and the principles governing 3D intercalation, essential for organogenesis, remain unclear. To explore how cells change shape and position during organogenesis, we studied tracheal system development in *Drosophila* embryos. This interconnected 3D network delivers oxygen to target tissues. The apical domains of tracheal cells face a central lumen, their lateral domains sustain cell-cell contacts, and their basal domains contact surrounding tissues. Subapical AJs, at the apical-lateral boundary, ensure intercellular adhesion. The embryonic tracheal tree is a compelling model of branched tubular epithelium that resembles structures in the mammalian kidneys, lungs, and mammary glands. Tracheal cells will collectively migrate in response to chemoattraction exerted by the ligand fibroblast growth factor (FGF) (Hayashi & Kondo, 2018). Notably, cellular rearrangements permanently maintain contacts between tracheal cells in the absence of cell division, and these rearrangements do not depend on Myo II’s contractile activity (Ochoa-Espinosa et al., 2017). We focused on the formation of the dorsal branches (DB), which depends exclusively on 3D intercalation mechanisms, in contrast with the dorsal trunk (DT), which is formed by cells that do not intercalate (Ribeiro et al., 2004). DB formation requires the progressive transformation of initial intercellular AJs (iAJs) into auto-cellular AJs (aAJs) (Fig. 1A) (Caussinus et al., 2008;

**Figure 1.**
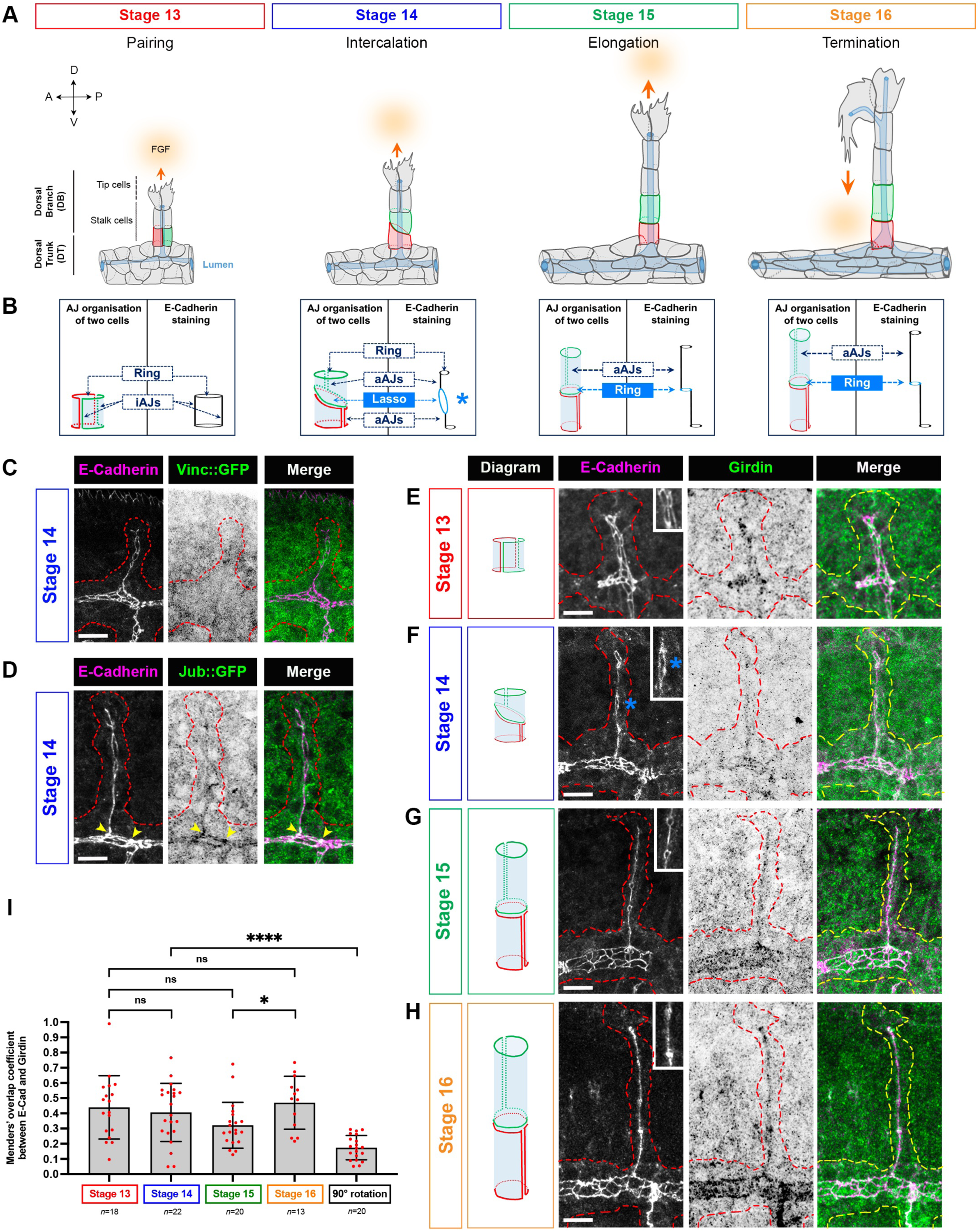
Girdin is enriched at the AJs of tracheal cells. (A) Stages in DB morphogenesis. The DB arises from the DT. DB cells, unlike DT cells, reorganise AJs and migrate toward the FGF ligand (orange oval) under pulling force from the tip cells (orange arrow). At st. 13 (pairing), green and red cells are paired on either side of the lumen (blue). At st. 14 (intercalation), they slide over each other. At st. 15 (elongation), intercalation is completed and the cells continue elongating until st. 16 (termination). Here, and in the remaining figures, anterior is left and dorsal is up. (B) Schematics of AJ organisation in green and red cells and the pattern of E-Cad staining at st. 13–16. The blue cylinder represents the lumen. At st. 13, cell pairs are side by side and are joined by iAJs, visible as two parallel lines of E-Cad staining. Top and bottom rings correspond to iAJs joining tracheal cells above and below the cell pair. At st. 14, dorsal migration of tip cells toward the FGF source exerts pulling forces that reposition paired cells and trigger intercalation. Each cell extends around the lumen to form aAJs. The iAJ–aAJ transition progresses as the green cell slides over the red one, visible as a line of E-Cadherin staining. The process of iAJs becoming aAJ results in a lasso-like organisation (blue star). At st. 15, the lasso transforms into a small ring of iAJs between the green and red cells. By st. 16, cell elongation extends the aAJ line. (C, D). Distributions of Vinculin (C) and Ajuba (D), detected by GFP antibody labelling in respective protein trap lines respectively (shown as inverted lookup tables (LUTs)), relative to E-Cad-labelled AJs in tracheal cells of the DT and DB. Yellow arrows and red dashed lines highlight the DT AJs and tracheal branches, respectively. (E–H) Schematics of the different stages of tracheal cell reorganisation (described in (B)) and E-Cad and Girdin immunofluorescence of the DT and DB at each stage. Girdin signal is shown as inverted LUTs. Red and yellow dashed lines highlight the tracheal structure. (I) Manders’ overlap coefficient (M2) calculated between Girdin and E-Cad at st. 13–16 and using clockwise-rotated Girdin signal at st. 14. Scale bars, 10 µm

Ribeiro et al., 2004). At the onset of DB formation, cells are aligned in a side-by-side arrangement, and all AJs are intercellular (iAJs). Then pulling forces, generated by the dorsal movement of the tip cells toward the source of FGF, cause the cells to shift their relative positions and intercalate (Caussinus et al., 2008). The iAJs are successively replaced by aAJs (Fig. 1A,B). Effective tracheal cell intercalation relies on the coupling of FGF-induced pulling forces with trafficking-driven reorganization of AJ-based adhesion. Through dynamic E-Cad internalisation and recycling, AJ disassembly is swiftly followed by adhesion reformation, ensuring precise and orderly cell intercalation (Shaye et al., 2008).

In this study, we show that the AJs of tracheal cells forming the DB lack Jub and Vinculin but do contain Girdin. Therefore, studying the formation of the DB’s 3D architecture provides an opportunity to uncover the function of Girdin, which may be masked by the Jub and Vinculin interactions in other AJs. The complete loss of Girdin, through removal of both maternal and zygotic contributions, causes many defects in epithelial morphogenesis, underscoring its critical developmental functions (Houssin et al., 2015). To dissect the cell-autonomous role of Girdin during tracheal development, we combined tracheal cell-specific RNA interference (RNAi) in loss-of-function *Girdin* zygotic mutants. In this sensitized genetic background, we reveal the important role of Girdin in regulating the pace of cell intercalation during DB morphogenesis. We show that Girdin and E-Cad functionally interact to ensure proper cell intercalation; however, Girdin does not regulate E-Cad endocytosis. In addition, Girdin organises the AJ-associated apical actin network, but not the interactions required for apical secretion of luminal products. Finally, by developing an innovative genetic tool that enables the selective *in vivo* depolymerization of the actin cytoskeleton specifically within tracheal cells, we demonstrate that the interaction between Girdin and AJ is independent of actin-mediated junctional tension. This finding thereby reinforces the role of Girdin as a key constituent of the AJ complex.

## Results

### Girdin localises to tracheal cell AJs

We aimed to identify factors that could modulate the Myo II-independent forces exerted on AJs during the intercalation and extension of DB tracheal cells. We assessed the localisations of the tension-sensitive proteins Vinculin and Jub during intercalation (st. 14), as they are involved in regulating the tension applied to AJs (Kale et al., 2018; Razzell et al., 2018). We detected green fluorescent protein (GFP)-fused Vinculin (Vinc::GFP; Maartens et al., 2016) at epidermal AJs (Fig. S1A); however, it was barely detectable in the AJs of DB (Fig. 1C and Fig. S1E ). Similarly, Jub (Ajuba::GFP, Flybase and BDSC) colocalised with E-Cad in epidermal cells (Fig. S1B) and to a lesser extent in tracheal cells forming the DT; however, it was barely detectable in the AJs of DB (Fig. 1D and Fig. S1D).

We next investigated if the cytoskeletal adaptor protein Girdin is expressed in the DBs using a specific anti-Girdin antibody (Houssin et al., 2015). Girdin is broadly distributed in epidermal cells in embryos of all stages (Houssin et al., 2015) and enriched at AJs (Fig. S1C). During morphogenesis, Girdin was expressed in all cells in the tracheal system, at higher levels in DT cells than in DB cells, and colocalised with E-Cad (Fig. 1E–H). We calculated Manders’ overlap coefficient (M2) to quantify the fraction of DB Girdin that overlapped with E-Cad. We found that 40% of Girdin is in the AJ domain, as the M2 value was approximately 0.4 (Fig. 1I), and this fraction remained constant throughout the formation of the DB. M2 values for Vinc::GFP and Jub::GFP remained around 0.1 from st. 13 to 16 (Fig. S1F, G). After clockwise rotation of the Girdin and GFP channels at st. 14, the M2 value dropped to 0.17 for Girdin (Fig. 1I) and was unchanged for Vinc::GFP and Jub::GFP (Fig. S1F, G). Altogether, these results indicate that M2 values 0.1 likely reflect random overlap from signal noise and further support that a large fraction of Girdin colocalises with AJs in DB tracheal cells while Vinculin and Jub are largely absent.

### Girdin contributes to DB morphogenesis by regulating the pace of cell intercalation

We explored the function of Girdin during DB formation by cell intercalation as illustrated in Fig. 1A and 1B. At stage 14, pairs of cells surrounding the lumen slide over each other in response to pulling forces from migrating tip cells. Intercalation starts at the branch base, as each cell wraps around the lumen to contact itself, forming aAJs (Ribeiro et al., 2004). This stage of sliding intercalation is characterised by a lasso-like organisation of the AJs (Fig. 1B,F). The cells then continue sliding until they are no longer side by side but are located one above the other. The lasso is then transformed into a small ring of iAJs, corresponding to the AJs between the two stacked cells. This process is repeated for the pair of cells immediately above them. By st. 15, most cells have completed their intercalation. The aAJs appear as a continuous E-Cad line with evenly spaced iAJ rings (Fig. 1B,G; Fig. 2A).

**Figure 2.**
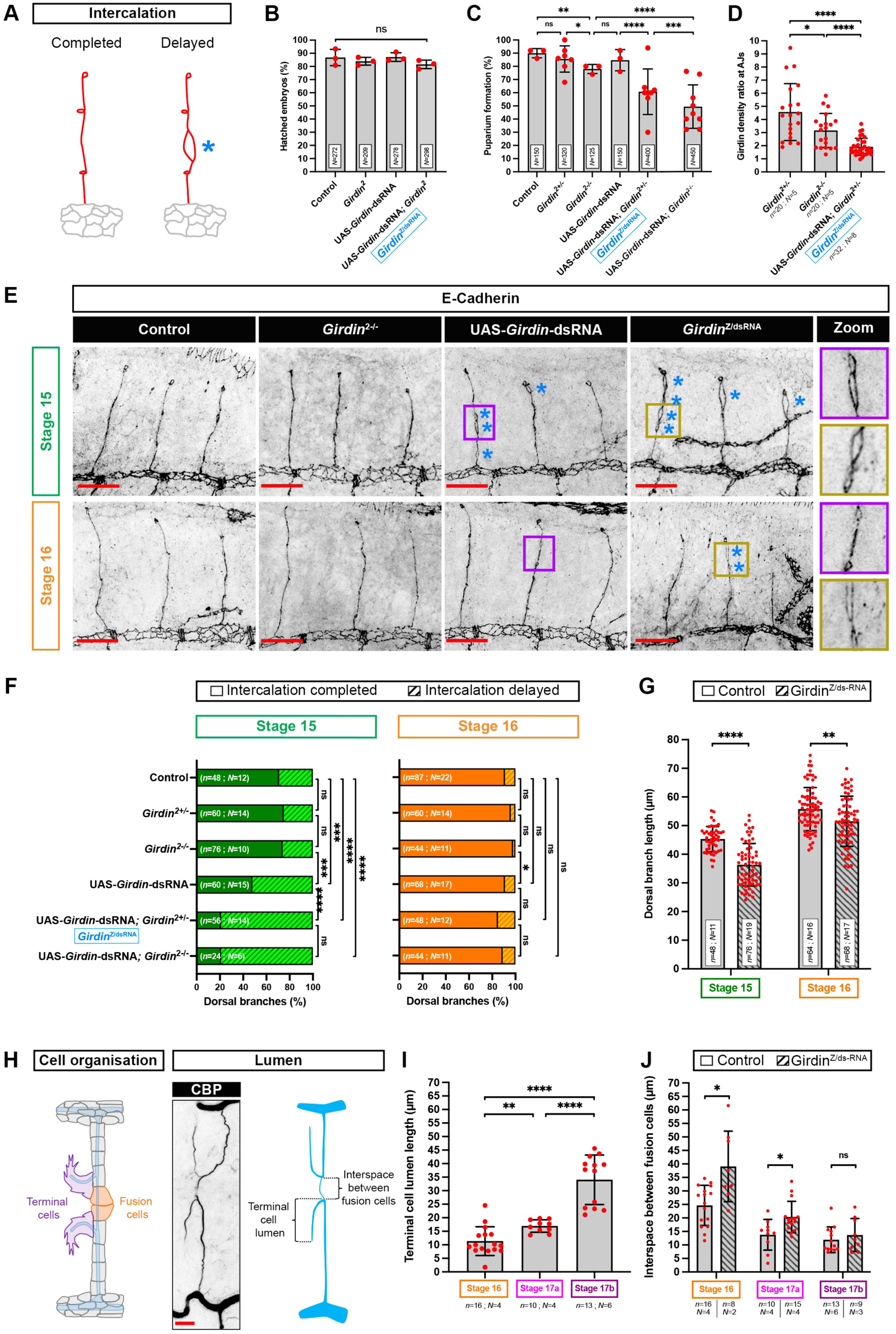
Girdin orchestrates intercalation timing during DB morphogenesis. (A) Schematic of AJ organisation at st. 15 with completed and delayed intercalation with a remaining lasso (blue star). (B, C) Successful embryo hatching (B) and puparium formation (C) in controls and *Girdin* mutants. Here and in all remaining figure panels except Fig. 3E-F, controls expressed the UAS-GFP::UTRN-ABD transgene under *btl*-GAL4 driver regulation. (D) Ratio of Girdin density at AJs in DB tracheal cells vs. epidermal cells in controls, *Girdin^2^* and *Girdin*^Z/dsRNA^ embryos. (E) E-Cad immunofluorescence to reveal the tracheal cell AJs in the st. 15 and 16 DBs of controls, *Girdin^2-/-^* zygotic mutants, UAS-*Girdin*-dsRNA expressing tracheal cells and *Girdin*^Z/dsRNA^ embryos (shown as inverted LUTs). Blue stars highlight residual lassos resulting from delayed intercalation. Zoomed-in views show intercalation at st. 15 and 16 in UAS-*Girdin*-dsRNA (purple) and *Girdin*^Z/dsRNA^ (brown) DBs. Scale bars, 20 µm. (F) Proportions of intercalation completed and delayed DBs at st. 15 and 16 in controls and *Girdin^2^*mutants with and without tracheal Girdin depletion. (G) DB lengths at st. 15 and 16 in the tracheal cells of controls and *Girdin*^Z/dsRNA^ embryos. (H) Schematic showing DB connections through fusion cells (orange) and terminal cell extensions (purple). Luminal continuity and terminal lumen extension, revealed by CBP staining, are shown in blue. (I) Lumen lengths in DB terminal cells at st. 16 and st. 17. Lumens that were 10–20 µm long and > 20 µm long were considered to be in st. 17a and 17b, respectively. (J) Interspaces between fusion cells at st. 16, st. 17a, and st. 17b in controls and *Girdin*^Z/dsRNA^ embryos.

Unambiguously distinguishing between the lasso formed at the end of sliding and the resulting ring can be difficult, depending on their orientations relative to the focal plane. We measured ring diameters at st. 15 and st. 16 in control DBs (see Materials and Methods and Fig. 1B). The mean diameter at st. 15 was 1.74 µm (± 0.51 µm), with a maximum of 2.74 µm; the ring diameter at st. 16 was 1.57 µm (± 0.55 µm), with a maximum of 3.18 µm. Therefore, any ring with a diameter > 3 µm at st. 15 was considered to be a lasso, signifying intercalation delay (Fig. 2A). In control embryos, > 70% of tracheal cells had completed intercalation by st. 15, with few residual lassos observed (Fig. 2E,F). By st. 16, almost all cells had completed intercalation (Fig. 2E,F), and the DBs had achieved their maximum length (∼56 µm; Fig. 2G) and reached the most dorsal part of the embryo. At this stage, when branch length is maximal, the two fusion cells from opposing branches are able to establish a new cellular contact, establishing continuity of the airway lumen (Fig. 2H; Kato et al., 2016).

To investigate Girdin function, we used a loss-of-function *Girdin^2^*zygotic mutant allele (*Girdin^2^*). Homozygotes for this allele do not die before the pupal stage (Fig. 2B,C) due to maternally contributed Girdin mRNA and protein (Houssin et al., 2015). As expected, no defects in DB formation were observed in *Girdin^2^* zygotic mutants at st. 15 compared to controls (Fig. 2E,F). Conversely, the complete loss of both maternal and zygotic Girdin function results in dramatic defects during embryogenesis, such as dorsal closure defects, impaired head morphogenesis, and loss of epithelial integrity (Houssin et al., 2015). These defects impact the entire embryo, which prevents us from determining if tracheal phenotypes are cell-autonomous or not. To overcome this, we combined dsRNA with the UAS/GAL4 system to specifically deplete Girdin from tracheal cells after placode invagination. Using the anti-Girdin antibody, we measured Girdin density ratio at the AJs of Girdin-depleted tracheal cells and in Girdin-expressing overlying epidermal cells (Fig. 2D and Fig S2A,B). This analysis revealed a lower tracheal-to-epidermis signal intensity ratio in Girdin-depleted embryos compared to control embryos, confirming the specific depletion of Girdin in tracheal cells. We also observed more residual lassos at st. 15 in Girdin-depleted DB cells, indicating incomplete intercalation (Fig. 2E,F). By st. 16 the lassos had largely disappeared upon Girdin depletion, indicating that intercalation was delayed but not supressed (Fig. 2E,F).

To further decrease Girdin expression in tracheal cells, we knocked-down *Girdin* in either heterozygous or homozygous *Girdin^2^* mutants. In both conditions, the Girdin density ratio at the AJs was reduced by more than half compared to heterozygous *Girdin^2^* mutants (Fig. 2D and data not shown). These genetic contexts did not induce embryonic lethality (Fig. 2B); instead, they increased larval lethality and precluded puparium formation (Fig. 2C). Larval lethality was further enhanced in *Girdin^2^*homozygotes depleted of Girdin. This further reduction in Girdin expression increased the number of lassos observed at st. 15 compared to Girdin depletion alone, with intercalation delays observed in 80% of DBs with depletion plus mutation, compared with 50% of DBs displaying lassos upon depletion alone (Fig. 2E,F). By st. 16, most lassos had disappeared and 90% of DBs had completed intercalation (Fig. 2E,F). As no significant differences were observed in the AJ Girdin levels or intercalation delays between the Girdin-depleted heterozygous and homozygous mutants, the remainder of our studies were performed using the heterozygous Girdin-depleted mutants, referred to as *Girdin*^Z/dsRNA^.

In *Girdin*^Z/dsRNA^ embryos at st. 15, which displayed delayed intercalation, DBs were significantly shorter than in controls (Fig. 2G). In control conditions, lumen continuity between two opposing DBs occurred at st. 16 and is distinguished by the extension of lumen markers, such as chitin-binding protein (CBP), which labels chitin fibrils between the two fusion cells (Fig. 2H) (Kato et al., 2016; Tonning et al., 2005). The final stage of embryogenesis (st. 17) is characterised by terminal cell elongation associated with growth of the intra-cytoplasmic lumen (Fig. 2H). As this stage lasts ∼5 hours, it was divided in two according to the length of the terminal cell lumen, with lumens measuring 10–20 µm and > 20 µm sorted into st. 17a and st. 17b, respectively (Fig. 2I). In *Girdin*^Z/dsRNA^ conditions, where DBs are shorter, the interspace between fusion cells of counterpart DBs was larger than in controls at st. 16, and airway lumen continuity was not yet achieved. The *Girdin*^Z/dsRNA^ embryos showed slightly larger interspaces between fusion cells than controls in st. 17a, but these interspaces progressively decreased by stage 17b, ultimately establishing luminal continuity between the DBs (Fig. 2J and Fig.S2C). Together, these results indicate that Girdin reduction does not prevent tracheal cell intercalation; however, it does decrease the speed of the process.

### Girdin interacts with E-Cad but does not regulate its endocytosis at the apical tracheal cell membrane

Because Girdin has been shown to contribute to E-Cad-mediated cell-cell adhesion in epithelial tissues, we explored whether the intercalation defects in *Girdin^2^* zygotic mutants could be enhanced by the reduction of E-Cad. *Drosophila* E-Cad is encoded by *shg*, and is preferentially required during cell rearrangement in the neurectoderm and other morphogenetically active epithelia (Tepass et al., 1996). In embryos homozygous for the *Girdin^2^* zygotic mutant allele (Fig. 2E,F) or the hypomorphic E-Cad allele *shg^119^* (Le Droguen et al., 2015; Tepass et al., 1996) no intercalation defects were observed at st. 15 and 16 compared to controls as revealed respectively by E-Cad staining and Polychaetoid (Pyd) staining, the *Drosophila* homolog of the junctional protein ZO-1 (Jung et al., 2006) (Fig. 3A,B). However, *shg^119-/-^*; *Girdin^2-/-^* double mutants displayed numerous residual lassos at st. 15 (Fig. 3A,B) as revealed by E-Cad staining. By st. 16, the number of residual lassos in the double mutants had decreased but remained significantly higher compared to single mutants (Fig. 3A,B). These results suggest that Girdin and E-Cad functionally interact to ensure timely cell intercalation.

**Figure 3.**
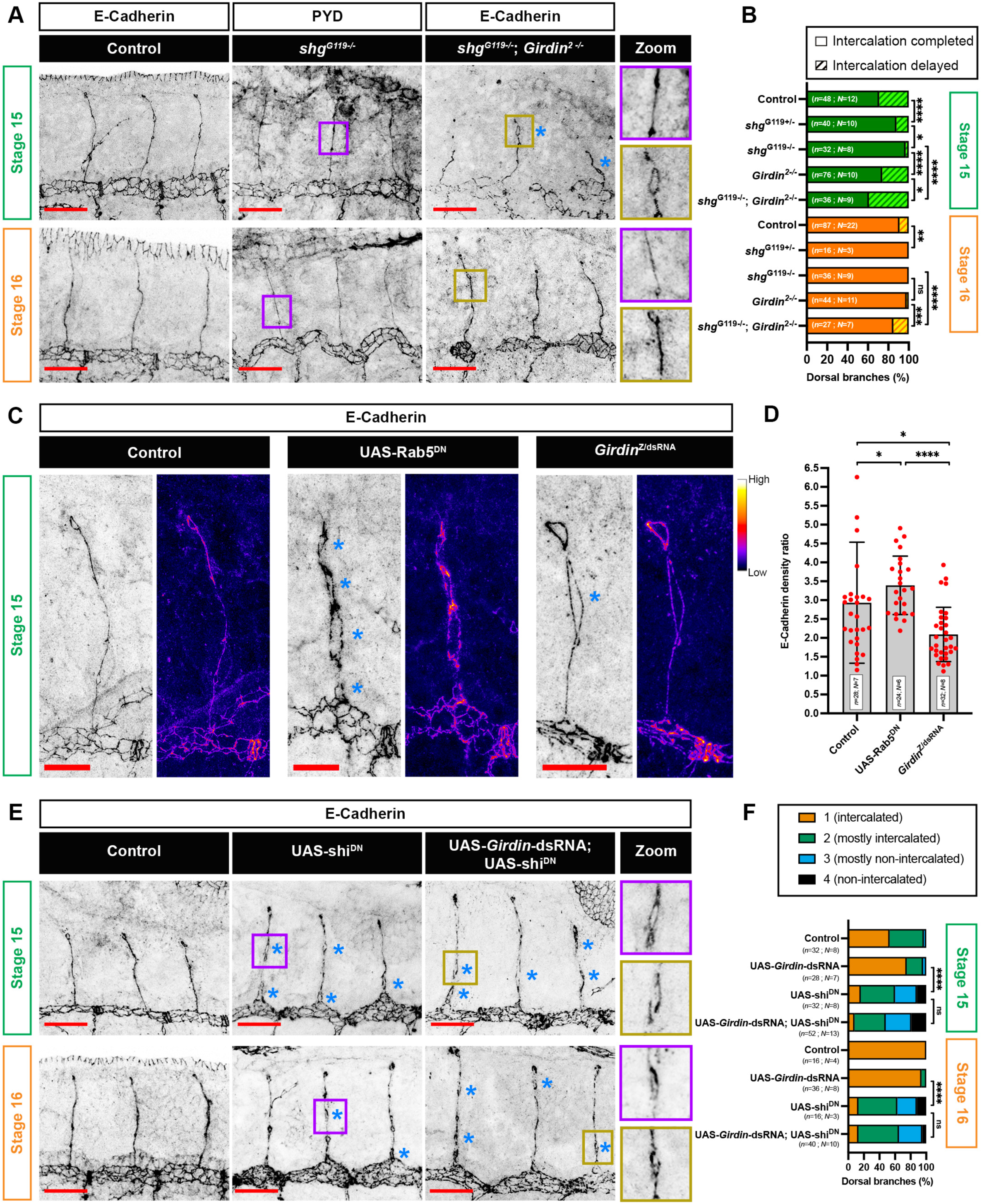
Girdin interacts with E-Cad independently of endocytosis. (A) Polychaetoid (Pyd) staining reveals tracheal cell organisation at st. 15 and 16 in *shg^G119-/-^*zygotic mutants and E-Cad staining in controls and *shg^G119-/-^*; *Girdin^2-/-^* zygotic double mutants (both as inverted LUTs). Residual lassos are indicated by blue stars. Purple and light brown boxes at st. 15 and st. 16 show the branch intercalation states in the *shg^G119-/-^* mutants and the *shg^G119-/-^*; *Girdin^2-/-^* zygotic double mutants, respectively. Scale bars, 20 µm. (B) Percentages of intercalation completed and delayed DBs at st. 15 and 16 in controls and the indicated mutants. Values for the *Girdin^2+/-^* zygotic mutants were those shown in Figure 2F. (C) E-Cad levels shown as invert LUTs and as Fire LUTs in control, Rab5^DN^-overexpressing, and *Girdin*^Z/dsRNA^ tracheal cells. Scale bars, 10 µm. (D) E-Cad density ratios at AJs in control, Rab5^DN^, and *Girdin*^Z/dsRNA^ DBs compared to epidermal cells. (E) E-Cad localisation at st. 15 and 16 reveals DB organisation in control and shi^DN^-expressing tracheal cells with or without Girdin depletion (inverted LUTs). The purple and light brown boxes show intercalation at each stage in shi^DN^ overexpressing cells with and without Girdin depletion, respectively. Blue stars mark residual lassos resulting from delayed intercalation. Scale bars, 20 µm. (F) Intercalation scores distribution at st. 15 and st. 16 in the control and shi^DN^ expressing tracheal cells with and without Girdin depletion. The controls contain the *btl*-GAL4 driver alone.

To further explore this interaction, we investigated the distribution of E-Cad in *Girdin^2^* zygotic mutants. In cultured mammalian cells, the Girdin ortholog CCDC88A interacts with dynamin 2, a GTPase that excises endocytic vesicles from the plasma membrane, promoting the endocytosis of certain cargos, including E-Cad. The internalisation of E-Cad is affected in *CCDC88A*-depleted Madin-Darby canine kidney cells (Weng et al., 2014). In tracheal cells, the E-Cad level is partly dependent on endocytosis (Shaye et al., 2008). Therefore, we examined whether Girdin regulates E-Cad through endocytosis, which could explain the intercalation delays observed in Girdin-depleted tracheal cells. To compare the E-Cad densities between different conditions, we measured the signal density and calculated the ratio between tracheal and epidermal levels. In control embryos, the density ratio was 2.9, indicating that E-Cad level was approximately three times higher in the tracheal system than epidermal cells (Fig. 3C,D and Fig.S3). As a positive control for impaired E-Cad internalisation, we expressed, in tracheal cells, a dominant-negative form of Rab5 (Rab5^DN^), which blocks endocytosis (Shaye et al., 2008). Rab5^DN^ expression reduced DB intercalation (Fig. 3C ; Shaye et al., 2008) by maintaining slightly but significantly higher E-Cad levels at the AJs compared to controls conditions, with a ratio of 3.4 (Fig. 3D and Fig. S3). We hypothesised that if Girdin regulates E-Cad endocytosis, E-Cad density ratio at the AJs should be higher in our *Girdin* mutant conditions. However, in *Girdin*^Z/dsRNA^ DBs, in which the Girdin level is significantly reduced (Fig. 2D), we measured a significant decrease in E-Cad density ratio of ∼2 (Fig. 3C,D and Fig. S3).

To confirm that Girdin does not play a role in E-Cad endocytosis by interacting with Dynamin 2 in tracheal cells, we expressed a dominant negative version of the dynamin 2 ortholog Shibire (shi^DN^), which partly prevents intercalation of DB cells by maintaining an abnormally high level of E-Cad (Shaye et al., 2008), and investigated whether reducing Girdin function would worsen this phenotype. As expected, expressing shi^DN^ resulted in an increase in remaining visible lassos at st. 15 and 16 (Fig. 3E). To quantify the severity of intercalation defects, we used a four-point scale, ranging from 1 (indicating a DB with complete intercalation) to 4 (indicating no intercalation) (Fig. 3F; Shaye et al., 2008). The mean score in embryos expressing shi^DN^ was 2.38 at both st. 15 and st. 16, compared with control scores of 1.50 and 1.00 at st. 15 and 16, respectively. Girdin depletion alone resulted in few residual lassos at st. 15 (Fig. 2E,F and Fig. 3F), with mean scores of 1.29 and 1.05 at st. 15 and st. 16, respectively. When shi^DN^-expressing embryos were depleted of Girdin, the number of remaining lassos was no higher than with shi^DN^ expression alone (Fig. 3E), with mean scores of 2.63 and 2.27 for st. 15 and st. 16, respectively (Fig. 3F). These results do not support a role for Girdin in regulating E-Cad endocytosis through an interaction with Dynamin 2 in tracheal cells.

### Girdin is required for actin enrichment at AJ

Previous studies have identified Girdin as a key player between the cadherin–catenin complex and the actin cytoskeleton (Enomoto et al., 2005; Houssin et al., 2015). Therefore, we tested whether Girdin contributes to the association of actin with AJs in tracheal cells by monitoring actin network distribution in controls and *Girdin*^Z/dsRNA^ embryos. Actin distribution was visualized by expressing the actin-binding domain (ABD) of the human F-actin binding protein Utrophin (UTRN) fused to GFP (GFP-UTRN::ABD; Rauzi et al., 2010) in tracheal cells only (Fig. 4A-C). The expression of this actin probe does not stabilise the actin network (Burkel et al., 2007) or interfere with fly viability when expressed in the tracheal system (Fig. 2B,C). In addition, the GFP-UTRN::ABD signal in tracheal cells colocalises with actin, as revealed by phalloidin staining (data not shown). At st. 15, *Girdin*^Z/dsRNA^ DBs had lower actin network levels than control DBs (Fig. 4A,B), and GFP-UTRN::ABD density was significantly reduced at AJs (Fig. 4C).

**Figure 4.**
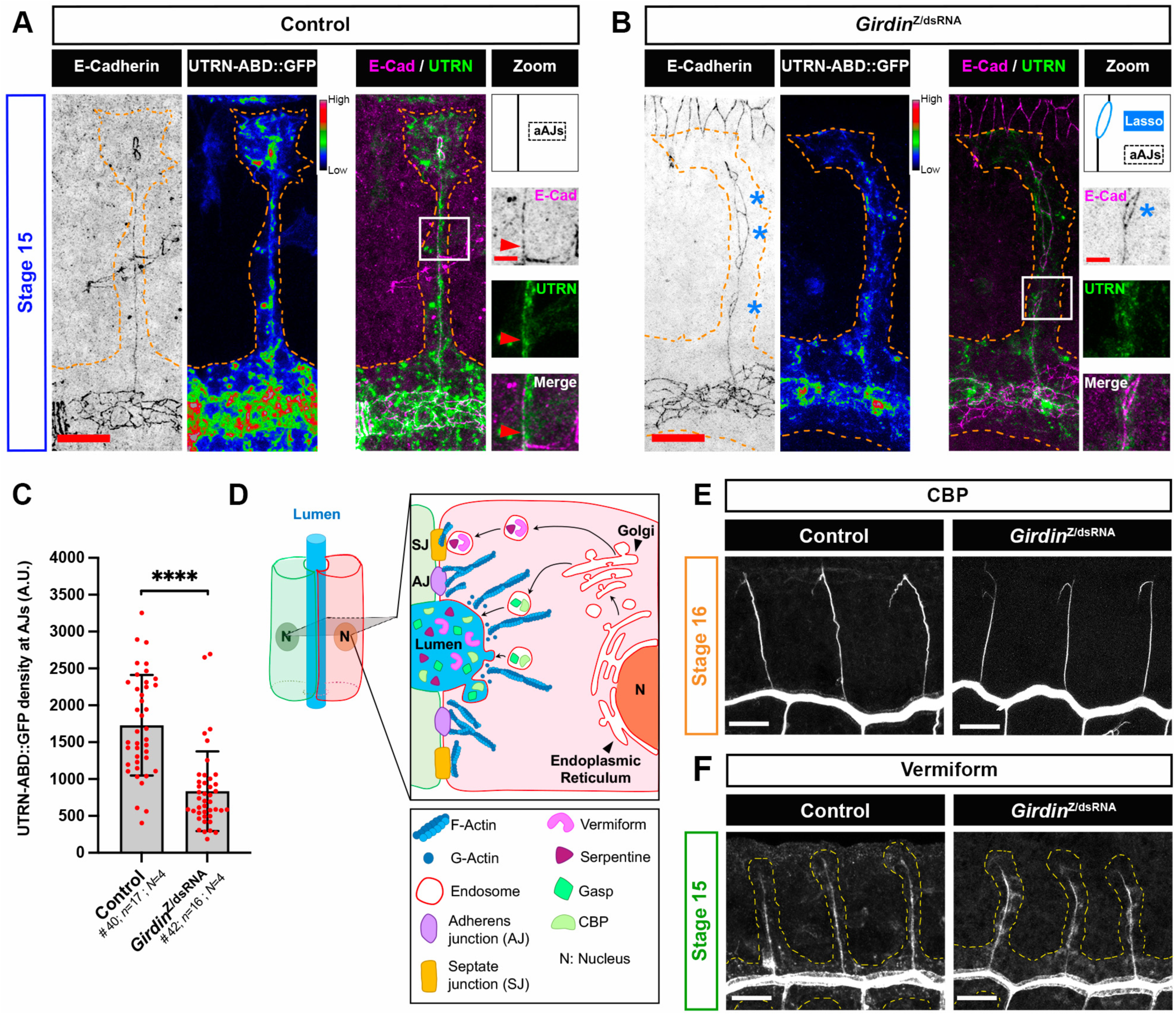
Girdin is required for actin enrichment at AJ but not for luminal product secretion. (A, B) UTRN-ABD::GFP localisation to E-Cad-labelled AJs in (A) control and (B) *Girdin*^Z/dsRNA^ DBs at st. 15. E-Cad and UTRN-ABD::GFP levels are shown in inverted and Rainbow RGB LUTs, respectively. Orange dotted lines delineate DB contours. Zoomed-in views (white rectangles) show E-Cad and UTRN-ABD::GFP at aAJs, appearing as a straight line in (A) and a line interrupted by a lasso in (B) (schematics). Red arrowheads indicate the AJs. Scale bars, 10 µm. (C) UTRN-ABD::GFP density quantifications at AJs in control and *Girdin*^Z/dsRNA^ DBs. (D) Schematic of luminal product secretion through the apical domain (Gasp, CBP) or *via* septate junctions (Serpentine, Vermiform). (E, F) CBP secretion at st. 16 (E) and Vermiform secretion at st. 15 (F) in the lumens of controls and *Girdin*^Z/dsRNA^ embryos. Scale bars, 20 µm.

These results indicate that in tracheal DBs, Girdin plays an important role in the association between the actin network and AJs. Another dense actin network lies beneath the entire apical surface of tracheal cells (Fig. 4D), which is distinct from the one associated with AJs and contributes to the secretion of products into the lumens of tracheal system (Massarwa et al., 2009). Therefore, we aimed to identify which of these two actin networks is compromised in the *Girdin*^Z/dsRNA^ embryos. As loss of the apical actin network is accompanied by defects in the secretion of many components destined to the lumen, such as Gasp, Piopio, and CBP (Massarwa et al., 2009), we studied CBP distributions in control and *Girdin*^Z/dsRNA^ DBs between st. 15 and st. 16. No defects or delays in secretion were observed (Fig. 4E); therefore, secretion is normal in *Girdin*^Z/dsRNA^ DBs, suggesting that the apical actin network is not altered or absent. Notably, not all products secreted into the lumen rely on the apical actin network; some, such as the chitin-binding deacetylases Vermiform and Serpentine, depend on the integrity of septate junctions (S. Wang et al., 2006; Fig. 4D). Vermiform deposition in the lumen was similar between control and *Girdin*^Z/dsRNA^ DBs (Fig. 4F), indicating that the Girdin depletion does not affect the integrity of septate junction.

### The role of Girdin in tracheal cells depends on α-Catenin

Work performed in *Drosophila* embryo epidermal cells demonstrated that destabilising the actin network by injecting latrunculin A reduces the level of E-Cad in AJs (Cavey et al., 2008; Letizia et al., 2019). Therefore, we focused on the function of actin at AJs and its correlation with E-Cad to characterize the intercalation delay phenotypes observed when Girdin levels are severely reduced. Girdin co-precipitates with α-Catenin and actin (Houssin et al., 2015); therefore, it could link the cadherin–catenin complex to actin filaments. We tested whether Girdin’s function in tracheal cells also requires α-Catenin. As α-Catenin is essential for the formation and maintenance of AJs (Sarpal et al., 2012), embryos carrying hypomorphic *α-Catenin* mutant alleles display phenotypes from gastrulation onwards. To circumvent this problem, we used a RNAi approach together with the UAS/GAL4 system to specifically deplete α-Catenin in the tracheal system. In control embryos, α-Catenin antibody labelling revealed a α-Catenin density ratio of ∼2.9 in tracheal cells compared to epidermal cells, consistent with the density ratio measured for E-Cad (compare Fig. 3D with Fig. 5B). In α-Catenin-depleted DBs, the ratio was significantly lower (∼1.6; Fig. 5A,B), indicating that α-Catenin RNAi efficiently reduced α-Catenin levels by a third in the tracheal system without affecting overall embryogenesis. At st. 15, > 70% of α-Catenin-depleted cells had completed intercalation, similar to control cells (Fig. 5C,D). At st. 16, most lassos had disappeared, and intercalation was complete in 95% of DBs (Fig. 5C,D). However, upon co-depletion of Girdin and α-Catenin, there were significantly more remaining lassos at st. 15 than with depletion of either protein alone (Fig. 5C,D). However, at st. 16, lassos were no longer visible (Fig. 5C,D). In addition, *Girdin*^Z/dsRNA^ DBs, the α-Catenin density ratio between tracheal and epidermal levels was also reduced (Fig. S4A, B). Altogether, these results demonstrate that α-Catenin and Girdin functionally interact to contribute to the pace of DB cell intercalation.

**Figure 5.**
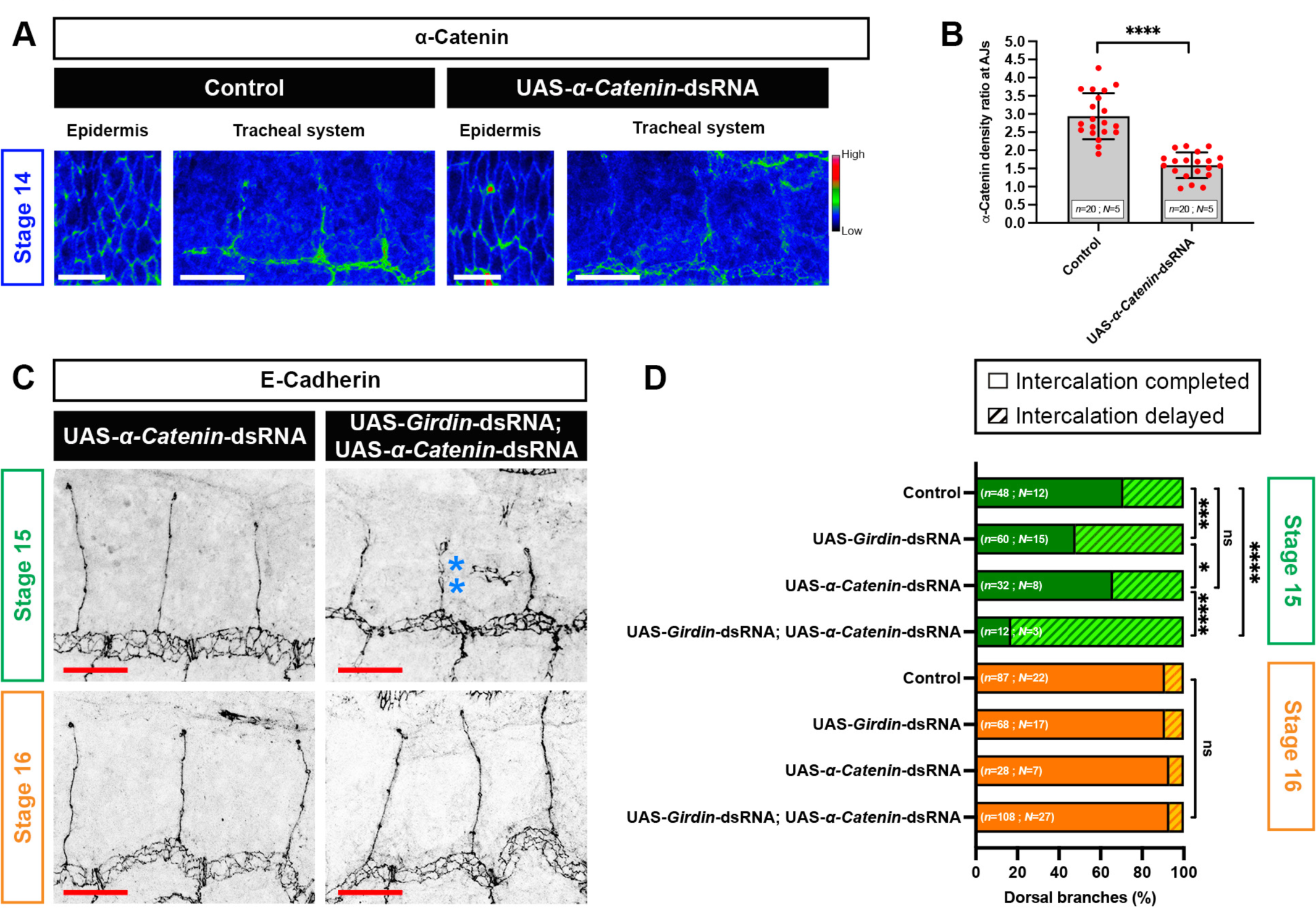
α-Catenin and Girdin promote efficient cell intercalation. (A) α-Catenin distribution at st. 14 in epidermal (scale bars, 10 µm) and tracheal cells (scale bars, 20 µm) in control and α-Catenin-depleted tracheal cells. Levels shown as RGB Rainbow LUTs. (B) Ratios of α-Catenin density at AJs between DB tracheal cells and epidermal cells in control and α-Catenin-depleted tracheal cells. (C) DB organisation revealed by E-Cad localisation at st. 15 and st. 16 in tracheal cells depleted of α-Catenin alone or together with Girdin. E-Cad shown as inverted LUT. Blue stars indicate residual lassos at st. 15. Scale bars, 20 µm. (D) Percentages of intercalated and intercalation-delayed DBs at st. 15 and st. 16 in control, Girdin-depleted, α-Catenin-depleted, and Girdin- and α-Catenin-depleted tracheal cells. Values for Girdin-depleted cells are identical to those shown in Figure 2D.

### Girdin is an integral component of AJs

Since Girdin was required for actin to associate with AJs, and its function depends on α-Catenin, we next tested whether the localisation of Girdin to AJs depends on its association with the actin network, as Vinculin does (le Duc et al., 2010). To do this, we developed a new genetic tool to depolymerise the actin cytoskeleton *in vivo*, only in tracheal cells. This tool is based on the expression of the mono-ADP-ribosyltransferase domain of *Salmonella enterica* SpvB (DeAct-SpvB) under the control of the UAS/GAL4 system. SpvB ADP-ribosylates actin monomers at a conserved arginine residue. These modified monomers can no longer be incorporated into new actin filaments, however the stability of the ribosylated monomers remains unknown. In all cases, SpvB activity leads to a net disassembly of all dynamic actin filaments (Fig. 6A); an outcome demonstrated both in cell culture and *in vivo* in multicellular model organisms, such as mouse and worms (Harterink et al., 2017).

**Figure 6.**
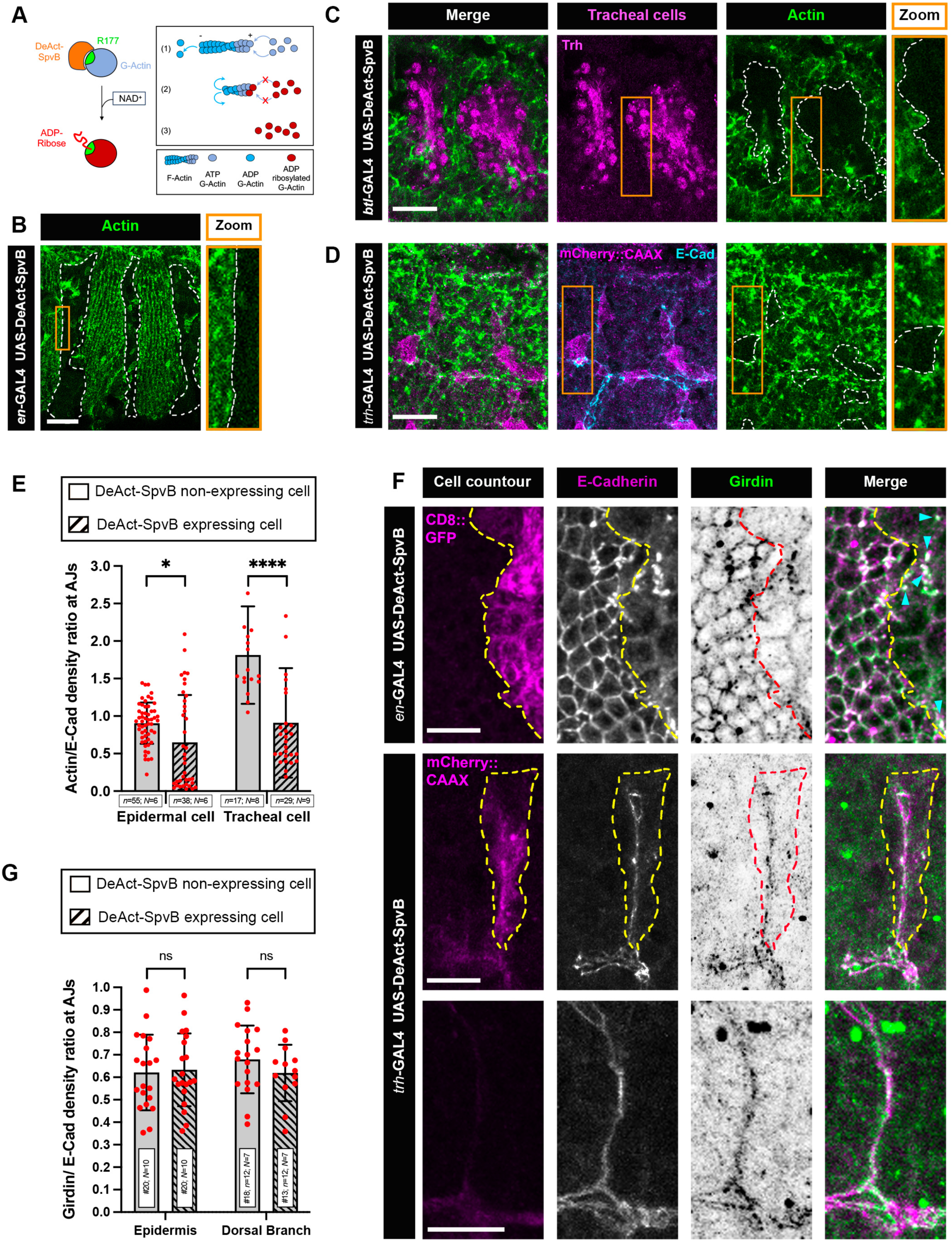
A novel genetic tool for cell-specific actin depolymerisation reveals Girdin as a key component of AJ. (A) Mechanism of action of the mono-ADP-ribosyl-transferase domain of SpvB (DeAct-SpvB). (1) DeAct-SpvB ADP-ribosylates actin monomers at a conserved arginine (R177), (2) inhibiting their polymerisation capacity and (3) causing net disassembly of dynamic actin filaments. (B–D) DeAct-SpvB expression in the posterior compartment (*en*-GAL4, st. 15) (B), in the tracheal system (*btl*-GAL4, st. 13) (C), and in a subset of tracheal cells (*trh*-GAL4, st. 14) (D). DeAct-SpvB effects were assessed by anti-actin labelling. Zooms (orange) span the GAL4 expression domains with white dotted lines marking boundaries. (C) Tracheal cells are identified by Trh expression and (D) mCherry-CAAX highlights GAL4-expressing tracheal cells. Cell positions within the tracheal system were determined according to their AJs, visualised by E-Cadherin. (E) Ratios of actin to E-Cad density in DeAct-SpvB expressing versus non-expressing cells, measured in both epidermal and DB cells. (F) Immunofluorescence of E-Cad and Girdin in epidermal and tracheal cells. DeAct-SpvB-expressing epidermal (*en*-GAL4) and tracheal (*trh*-GAL4) cells are identified by CD8::GFP and mCherry-CAAX, at st 10 and st. 15 respectively. DB without CAAX-positive clones (bottom panel) serves as a control for Girdin distribution. Dotted lines mark expression boundaries and blue arrowheads indicate regions where E-Cad remains high in DeAct-SpvB-expressing epidermal cells. Girdin is shown as inverted LUTs. (G) Ratios of Girdin to E-Cad density at AJs in DeAct-SpvB expressing versus non-expressing cells, measured in both epidermal and DB cells. Scale bars, 20 µm (B-D) and 10 µm (F).

To determine whether DeAct-SpvB expression could also induce actin depolymerisation in *Drosophila* cells, we used the *en*-GAL4 driver line, which expresses the GAL4 factor in all posterior compartments of embryonic segments. Significant depolymerisation of the actin network was observed in these cells only (Fig. 6B). Therefore, the GAL4-driven expression of DeAct-SpvB provides a powerful new tool for cell-specific perturbation of the actin cytoskeleton in *Drosophila*.

Using the pan-tracheal *breathless* (*btl*)-GAL4 line, we first determined that DeAct-SpvB overexpression induced strong actin depolymerisation only in tracheal cells (Fig. 6C and Fig. S5A). However, the effects were too deleterious for tracheal morphogenesis: the placodes invaginated but the branches did not bud (Fig. 6C). To restrict actin depolymerisation to a limited number of cells within the DB, we controlled DeAct-SpvB expression with the *trachealess* (*trh*)-GAL4 line to induce a so-called ‘mosaic’ expression pattern in which GAL4 is randomly expressed in only a subset of tracheal cells within a population of control cells (Kondo & Hayashi, 2013). The *trh*-GAL4 line was combined with a UAS-CAAX::mCherry line to identify GAL4-positive cells *via* mCherry expression. In this context, DeAct-SpvB expression induced complete actin depolymerisation in mCherry-positive cells but not in control mCherry-negative cells (Fig. 6D). Using *trh*-GAL4 and *en*-GAL4 lines, we then examined Girdin distribution in actin-depleted tracheal cells and epidermal cells, respectively (Fig. 6F). Consistent with previous studies involving latrunculin A injection in embryos (Cavey et al., 2008; Letizia et al., 2019), DeAct-SpvB–induced actin depolymerization led to a reduction in E-Cad levels in both tracheal and epidermal cells. To rule out the possibility that the residual E-Cadherin signal results from incomplete actin depolymerisation, we measured the actin/E-Cad density ratio in DeAct-SpvB–expressing tracheal and epidermal cells compared to controls. In both cell types, the ratio was reduced, indicating that actin density decreases more strongly than E-Cadherin density (Fig. 6E). These results confirm that DeAct-SpvB efficiently depolymerises the actin network *in vivo* in *Drosophila* cells. Given the established physical and functional association between Girdin and E-Cad (Houssin et al., 2015), a corresponding decrease in Girdin levels was anticipated under these conditions. Instead of measuring only Girdin density ratio at AJ, we quantified the ratio of Girdin-to-E-Cad signal intensity ratio in DeAct-SpvB–expressing tracheal and epidermal cells, and compared it to that of control cells. Although E-Cad levels decrease after DeAct-SpvB-mediated F-actin depletion in the trachea, Girdin proportion at the E-Cad domain remains unchanged in both cell types (Fig. 6F, G). To further support our conclusions, we calculated the M2 Manders’ coefficient to quantify the fraction of Girdin overlapping with E-Cad in actin-depolymerised tracheal cells (Fig. S5B). The M2 value remained at 0.4, similar to controls (Fig. 1I). These results confirm that Girdin localisation at AJs is independent of actin–AJ association, supporting its role as an important AJ component. Nevertheless, Girdin functions as a key linker between the junction domain and actin cytoskeleton (Fig. 4A,B). This distinction allowed us to assess Girdin recruitment to AJs independently of actin-mediated membrane tension, unlike Vinculin.

## Discussion

A key enduring question in morphogenesis is how cells connect AJs to the cytoskeleton. This linkage must be dynamic and force-responsive to accommodate cell rearrangements and shape changes. Previous studies have established that AJ–actin linkage involves complex multivalent interactions that ensure proper timing of development (Pinheiro & Bellaïche, 2018). However, the mechanism underlying this process in tissues where Myo II and Jub are not required, and where Vinculin shows no functional contribution, remains unclear. Our results reveal that Girdin is important for this alternative linkage mechanism. By introducing genetic changes that did not impair overall embryonic development, we revealed the importance of Girdin in controlling the rate of cell intercalation in DBs of the tracheal system. *Girdin*^Z/dsRNA^ embryos revealed that Girdin controls the level of E-Cad at AJs independently of E-Cad endocytosis. In addition, Girdin’s function requires α-Catenin, and Girdin is involved in the apical enrichment of the actin network. Both contribute to the enrichment of the actin network at AJs without affecting the actin network involved in the secretion of luminal products from the apical surface.

By developing a unique genetic tool that allows *in vivo* actin depolymerisation, we demonstrate that the interaction between Girdin and AJ does not depend on actin network, highlighting Girdin as an important component of the AJ complex.

We discovered that Girdin participates in cell intercalation by contributing its timing during AJ reorganisation. When Girdin levels were reduced, intercalation was delayed. This was made apparent by the presence of numerous lassos formed by AJs that are still being reorganised at st. 15 when intercalation is normally complete. However, no phenotype was observed in the DT of Girdin-depleted embryos despite its high levels in this tissue. Notably, DT morphogenesis does not require cell intercalation (Ribeiro et al., 2004), which strongly suggests that Girdin’s main function in the trachea is to support cell intercalation. In a wound healing assay using Vero cells, CCDC88A depletion led to disorganisation of the actin network and reduced cell migration (Enomoto et al., 2005). This suggests that the delay in intercalation could be due to slower cell migration caused by decreased actin levels, especially at the AJs. We have not measured the speed of tracheal cell migration in *Girdin* mutants, but we suggest that the intercalation rate is a reliable proxy for migration velocity.

Although Girdin depletion delayed intercalation and thereby hindered the timely establishment of luminal continuity between opposing DBs (Fig. 2J, S2C), the viability of the hatched mutant embryos remained comparable to that of the controls (Fig. 2B). However, in *Girdin*^Z/dsRNA^ embryos, the tracheal cells remained hypomorphic for Girdin, with low levels still detected at the AJs (Fig. 2D), likely due to strong maternal contributions of proteins and mRNA. However, these were the only conditions that allowed Girdin function in the tracheal system to be observed without compromising overall embryonic integrity. The delay, rather than the hold, in intercalation could also be attributed to functional redundancy among multiple molecules that link AJs to the actin network. Given the critical need for cells to change shape and position within a tissue while preserving tissue integrity, such redundancy would not be surprising. Ensuring a robust connection between AJs and the actin network likely requires a diverse set of adapter proteins, allowing junctions to maintain both strength and flexibility.

The observed E-Cad reduction levels in our *Girdin*^Z/dsRNA^ conditions were likely a result of the reduction in actin at the AJs; as previously demonstrated in epidermal cells, latrunculin A induced actin filament depolymerisation reduces E-Cad levels in AJs (Cavey et al., 2008; Letizia et al., 2019). Girdin has been shown to contribute to actin organisation in various cell types. For example, in the skin cancer cell line A431, CCDC88A binds to α- and β-catenin, and its depletion leads to the disorganisation of the actin cytoskeleton (X. Wang et al., 2018). Studies in Vero cells show that CCDC88A plays a role in the regulation of actin cytoskeleton remodelling. In addition, CCDC88A C-terminus can bind to F-actin (Enomoto et al., 2005), and electron microscopy and immunogold labelling of Vero cells revealed that CCDC88A molecules localise with actin filaments, and higher magnifications demonstrated extensive colocalisation of CCDC88A to the junctions between the actin filaments. Importantly, our work indicates that the actin network required in tracheal cells for luminal secretion is not affected in *Girdin*^Z/dsRNA^ embryos.

The decrease, rather than increase, in E-Cad levels in our *Girdin*^Z/dsRNA^ embryos does not support a role for Girdin in the regulation of E-Cad endocytosis. This is further supported by the absence of enhanced intercalation defects upon DN dynamin 2 expression when Girdin levels were reduced. Furthermore, the intercalation delays observed in the *Girdin*^Z/dsRNA^ DBs cannot be explained by an impaired ability to reorganise AJs due to reduced E-Cad levels; indeed, intercalation remained unaffected in the hypomorphic *shg^G119-/-^* mutants. These delays are likely due to Girdin’s additional role in linking the actin network to AJ complexes, which distinguishes the defects from *shg^G119-/-^* single mutants.

Cadherin-based AJs mediate cell–cell adhesion, regulate cellular rearrangements (e.g., cell intercalation) and serve as anchoring sites for the actin cytoskeleton. β-catenin binds directly to the cytoplasmic tail of E-Cad, while α-Catenin interacts with β-catenin. In turn, actin filaments bind to α-Catenin. However, α-Catenin does not provide a stable physical link to actin filaments. Importantly, α-Catenin is not the sole molecule modulating the connection between the AJ complex and the actin cytoskeleton. Other linker proteins also fulfil this role, conferring distinct mechanical properties to AJs depending on the specific cellular context. Our work reveals that Girdin, in epidermal and tracheal cells, is an integral component of AJs, acting as a key linker between the AJ complex and the actin cytoskeleton. Girdin contributes to the stability of the AJ complex, as reduced levels of E-Cad are observed in its absence. As an integral component, Girdin’s association with the AJ complex is independent of the actin cytoskeleton, unlike other α-Catenin binding partners such as Vinculin and Jub (le Duc et al., 2010; Sheppard et al., 2023).

During morphogenesis, AJ remodelling involves multiple mechanisms targeting either the cadherin-based complex itself or its interaction with the actin cytoskeleton. Both types of mechanism rely on diverse set of linker proteins. Understanding how these AJ-actin linker proteins are recruited to AJs is essential for elucidating how cells dynamically change their shape and position. To investigate their respective inputs, we have developed the first *in vivo* tool in *Drosophila* that allows actin network depolymerization with spatial and temporal precision. This approach enables the discrimination between linker proteins that associate with the cadherin complex independently of actin and those that require actin binding to localise to AJs. This will enhance our understanding of how AJ-associated proteins modulate cellular rearrangements during tissue morphogenesis.

## Materials and methods

### D. melanogaster strains

Descriptions of most genetic elements can be found at http://flybase.net (FlyBase Consortium, 1999). *D. melanogaster* stocks and crosses were kept under standard conditions. The following mutations were used: *shg^G119^* (a gift from J. Zallen; Fox et al., 2005) and *Girdin^2^* (Houssin et al., 2015). CyO-*wg*-lacZ and TM6-*Ubx*-lacZ blue balancers were used to identify homozygous embryos. The GAL4 system (Brand & Perrimon, 1993) was used for misexpression experiments. The pan-tracheal *btl*-GAL4 driver (Shiga et al., 1996) was used at 18°C either in combination with UAS-shi^K44A(3-10)^ as shi^DN^ alone (Moline et al., 1999) or in combination with UAS-*Girdin*-dsRNA (BDSC#67960); at 25°C with UAS-GFP::UTRN-ABD (Rauzi et al., 2010) and UAS-DeAct-SpvB (this work); or at 29°C with UAS*–α-Catenin*–dsRNA (BDSC #33430), UAS-Rab5^DN^ ^(S43N)^ (Entchev et al., 2000), and UAS-*Girdin*-dsRNA alone or associated with the *Girdin^2^* mutant allele. The *trh*-GAL4, UAS-mCherry::CAAX driver (Kondo & Hayashi, 2013) and the *en*-GAL4; UAS-CD8::GFP driver (Brodu et al., 2004) were used at 25°C in combination with UAS-DeAct-SpvB. Vinc::GFP (Maartens et al., 2016) and Jub::GFP (BDSC#56806) were used to follow their respective distribution in epidermal and tracheal cells. Experiments were performed on fixed embryos, followed by immunolabelling. Fluorescently tagged proteins were detected using antibodies against the corresponding fluorescent tags.

### Plasmid constructs

The mono-ADP-ribosyl-transferase domain of *S. enterica* SpvB (DeAct-SpvB, amino acids 375–591) was cloned into the pUASz1.0 vector (Drosophila Genomics Resource Center #1431). The transgene was inserted by PhiC31-mediated transgenesis at the P{CaryP} attP2 docking site at 89E11 by BestGene Inc. The DNA construct was verified by sequencing. Full plasmid sequences or maps are available upon request.

### Immunohistochemistry and antibodies

Embryos were staged as described in Campos-Ortega & Hartenstein (1985) and fixed and stained following standard protocols (Le Droguen et al., 2015). Immunostaining was performed on embryos fixed in 4% formaldehyde for 20 min, or for 10 min for anti-E-Cad staining. Primary antibodies were as described in (Le Droguen et al., 2015) with the following additions: anti-α-Catenin (1/20; Developmental Studies Hybridoma Bank (DSHB)), anti-Girdin (1/500; Houssin et al., 2015), anti-Trachealess (1/100; Brodu & Casanova, 2006), anti-actin (1/500; MP Biomedicals clone C4), anti-Polychaetoid (Pyd) (1/150; DSHB), anti-Green Fluorescent Protein (GFP) (1/500; Aves Lab), anti-Red Fluorescent Protein (RFP) (1/500; Clontech), anti-Vermiform (1/500; S. Wang et al., 2006), anti-β-galactosidase (1/1000; Cappel), and CBP-647 (1/200; gift from N. Marti and J. Casanova). Secondary antibodies conjugated with Cy3 or Cy5 (The Jackson Laboratory) or Alexa Fluor 488, 546, or 647 (Life Technologies) were used at 1/200.

### Image acquisition

Images of fixed tissues were acquired by confocal microscopy on a Zeiss LSM 700 confocal microscope or a Zeiss LSM 980 Airyscan FLIM confocal microscope with a super-resolution module for actin distribution, both equipped with 488 nm, 561 nm, and 639 nm excitation laser lines and a 63× oil immersion objective (NA 1.4). Images were acquired with ZEN confocal software (Zeiss, version 3.5) and processed using Fiji (Schindelin et al., 2012) and Photoshop (Adobe Photosystems, version 25). Images were projections of a few consecutive confocal sections spaced 1 µm apart on the LSM 700 microscope and 0.3 µm apart on the LSM 980 Airyscan FLIM microscope. To compare values between sections, we integrated the decrease in signal intensity along the Z-axis on all images, as described in Brodu et al. (2010).

### Image analyses

Analyses were performed on the same four dorsal central branches (abdominal segments 1–4) to ensure reproducibility, and processed in Excel (Microsoft, version 16). The reference line used as a control expressed the UAS-GFP::UTRN-ABD transgene under the regulation of the *btl*-GAL4 driver.

### Quantification of actin density at AJs

Images were acquired with the Zeiss LSM 980 Airyscan FLIM confocal microscope using the same settings for control and *Girdin*^Z/dsRNA^-expressing mutant DBs. Actin levels were assessed by measuring GFP::UTRN-ABD expression in both conditions in projections of 2–5 sections. To do this, we developed a three-step macro in Fiji: 1) a region of interest (ROI) was selected within one tracheal cell of the DB; 2) in this ROI, AJs were automatically delimited based on E-Cad staining and defined as reference points, with AJ domains spanning 0–300 nm from the reference point; and 3) the density of the GFP signal was quantified from this reference point to the basal membrane along the entire height of the ROI. As ROIs had different heights, the data were normalised per unit area.

### Colocalisation between E-Cad and Girdin

Colocalisation between E-Cad and Girdin staining was measured by obtaining M2 Manders’ coefficients using the JaCop plug-in in Fiji. Within a ROI delimited to a DB outline, each channel was split and converted to a 16-bit image to implement the JaCop plug-in. In JaCop, manual thresholding was performed to remove background noise from the analysis, except for the Girdin signal. Randomisation was also performed. The M2 coefficient was calculated corresponding to the fraction of the Girdin signal in areas containing the E-Cad signal, with 0 and 1 indicating no colocalization and complete colocalisation, respectively.

### Relative levels of E-Cad, Girdin, and α-Catenin in different *Girdin* mutants

Ratios were calculated between the signal densities in the AJs of DBs and the AJs of epidermal cells. AJs were delimited based on the E-Cad signal, and signal densities were measured using Fiji’s mean value (the sum of the pixel intensity in the ROI/the number of pixels in the ROI) using projections of 2–3 sections. Ratios were calculated between the signal densities of E-Cad, Girdin, and α-Catenin in the AJs of DBs and the AJs of epidermal cells in control, *Girdin^2^*, Girdin-depleted, *Girdin*^Z/dsRNA^, Rab5^DN^, and α-Catenin-depleted DBs.

### Relative levels of Girdin and E-Cad in actin-depleted cells

For the DB of st. 14 and 15 embryos, ratios were calculated between the signal densities of Girdin and E-Cad in the AJs between the DeAct-SpvB expressing tracheal cells, identified by mCherry-CAXX expression, and control tracheal cells. For epidermal cells of st. 10 embryos, similar ratios were calculated between DeAct-SpvB expressing epidermal cells, revealed by CD8::GFP expression, and adjacent control epidermal cells.

### Quantification of intercalation delays

To discriminate between lassos formed during intercalation and rings resulting from completed intercalation, > 40 rings (at st. 15 and 16) were measured under control conditions based on E-Cad staining. Rings > 3 µm in diameter were considered to be lassos, as this value corresponds to twice the average ring diameter, helping to avoid confusion between rings and lassos.

### Quantification of the viability of *Girdin* mutants at 29°C

Embryos were collected at 29°C on an agarose plate after overnight egg laying and counted under a binocular microscope (Zeiss STEMI 2000). The hatched larvae were counted after 24–48 hours, and the difference from the original count revealed the lethality before and during hatching. Groups of 50 larvae were collected from these plates and placed in tubes at 29°C. Pupae were counted after 5–8 days, and the number was subtracted from 50 to measure lethality during the larval stages. At least triplicate counts were performed for each condition.

### DB length measurements

At st. 15–16, straight lines were overlaid on the E-Cad signal from the base of each DB to its tip in control and *Girdin*^Z/dsRNA^ embryos to measure the DB lengths in µm.

### Analysis of fusion delay between counterpart DBs

We measured the interspace between fusion cells as a proxy of luminal branch fusion in control and *Girdin*^Z/dsRNA^ DBs at st. 16–17. These stages were determined by the length of the lumen in the terminal cell, revealed by CBP staining. Because the duration of st. 17 is substantial (5 hours), it was divided into two parts according to the length of the terminal cell lumen (see Fig. 2H,I). Terminal cells with 10–20 µm lumens were considered to be in st. 17a, while those with lumens > 20 µm were considered to be in st. 17b.

### Intercalation scoring

For DBs, intercalation was measured in st. 15 or 16 embryos using a four-point scale, as defined in Shaye et al., 2008: 1, intercalated DBs; 2, DBs with primarily aAJs (mostly intercalated); 3, DBs with primarily iAJs (mostly non-intercalated); and 4, DBs with only iAJs (non-intercalated). We calculated the mean DB score for each condition.

### Statistical analyses

GraphPad Prism 10 (GraphPad Software Inc., USA) was used to create graphs and perform statistical tests. Data represent the mean ± standard deviation (s.d.). The figure legends indicate the numbers of ROIs (“#”), DBs (“*n*”), and embryos (“*N*”), except for Fig. 2B and 2C, in which “*N*” corresponds to individuals. “*n*” and “*N*” values included at least four embryos per genotype. For means, we used the two-tailed non-parametric Mann–Whitney test when the data were not normally distributed (Figs 1I, 2D, 2G, 2G2I–J, 3D, 4C, 5B, 6E and 6G); for proportional data, a chi-squared test was performed (Figs 2B-C, 2F, 3B, 3F, 5D). Statistical significance is represented in the figures as follows: ns, not significant; *, *p* < 0.05; **, *p* < 0.01; ***, *p* < 0.001; ****, *p* < 0.0001.

## Acknowledgements

We acknowledge the ImagoSeine core facility at the Institut Jacques Monod, a member of France-BioImaging (ANR-10-INBS-04) and IBiSA, with the support of Labex “Who Am I” Inserm Plan Cancer, Region Ile-de-France, and Fondation Bettencourt Schueller.

We are grateful to Jordi Casanova for reagents and *D. melanogaster* strains and Martin Harterink and Casper Hoogenraad for the DeAct-SpvB plasmid. We also acknowledge the Bloomington *Drosophila* Stock Center for fly stocks and the DSHB for monoclonal antibodies.

We thank Jordi Casanova and Jean-Antoine Lepesant for discussion and critical comments on the manuscript and High-Fidelity Science Communications for manuscript editing.

## Data and resource availability

All relevant data and details of resources can be found within the article and its supplementary information.

## Author contributions

S.C. conducted most of the experiments together with V. B., P. L. and A. G. helped with data interpretation. V. B. designed the study and wrote the manuscript. All authors were involved in the final stages of manuscript writing.

Conceptualisation: V.B., A.G.

Methodology: V.B., S.C.

Software: V.B., S.C.

Validation: V.B., S.C., A.G., P.L.

Formal analysis: V.B., S.C., A.G.

Investigation: V.B., S.C., A.G.

Resources: V.B., A.G.

Data curation: S.C.

Writing - original draft: V.B.

Writing - review & editing: V.B., S.C., A.G., P.L.

Visualization: V.B., S.C., A.G.

Supervision: V.B., A.G.

Project administration: V.B., A.G.

Funding acquisition: V.B., A.G.

