## Supplementary data for "Girdin controls the pace of 3D tracheal cell intercalation by coupling adherens junctions to the actin cytoskeleton in *Drosophila*"

Fig. S1 Carvalho et al.

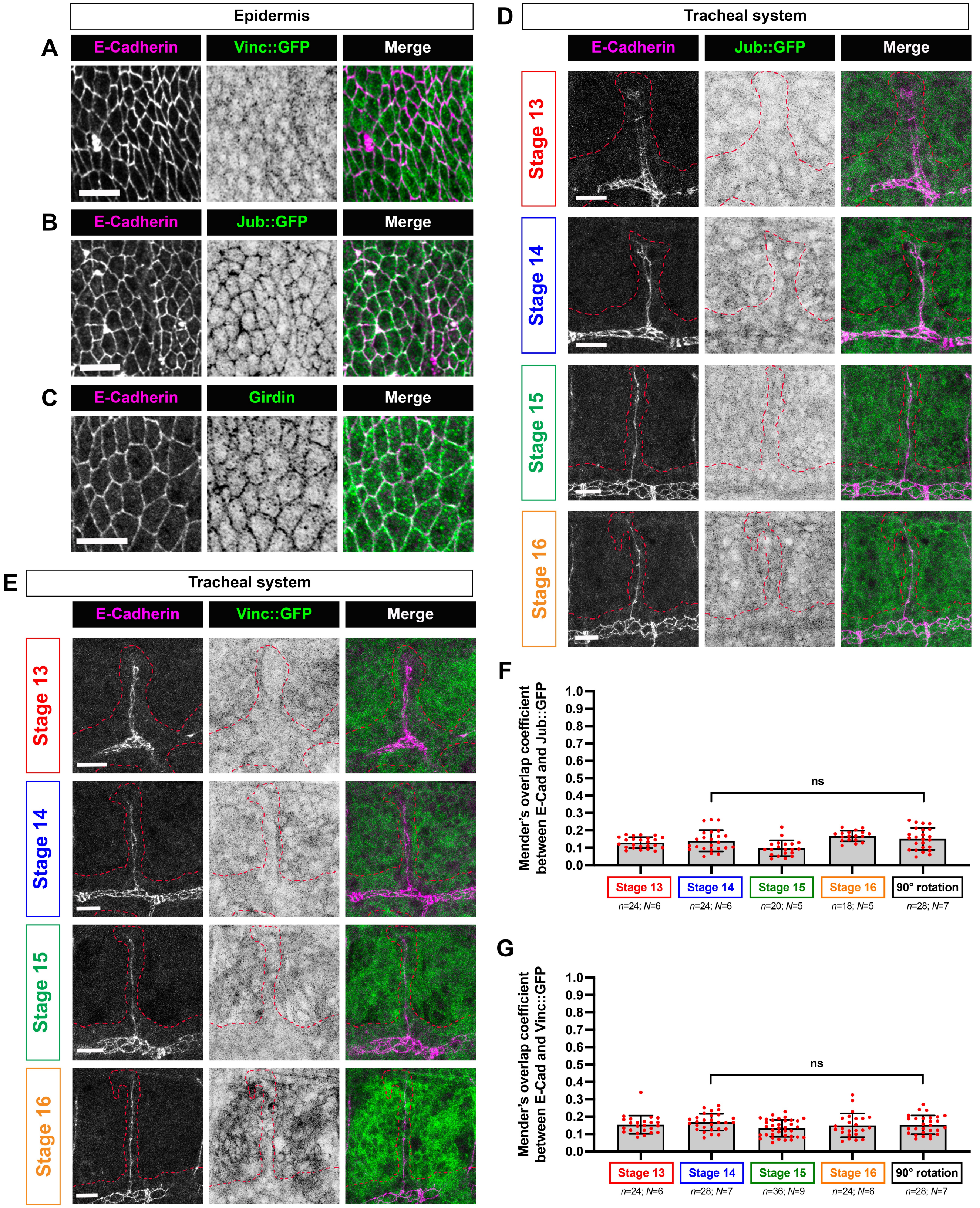

**Figure S1. Vinculin, Jub, and Girdin distributions in epidermal and tracheal cells**

(A-C) Distributions of Vinc::GFP (A) and Jub::GFP (B) and Girdin (C) relative to E-Cad-labelled AJs in epidermal cells. The Vinc::GFP and Jub::GFP distributions represent projections of two consecutive confocal sections, whereas Girdin distributions represent single confocal section. (D, E) Distributions of Jub::GFP (D) and Vinc::GFP (E) relative to E-Cad-labelled AJs in tracheal cells of the DT and DB from st. 13 to 16. GFP signal is shown as inverted LUTs. Red dashed lines highlight the tracheal structure. (F, G) Manders' overlap coefficient (M2) between Jub::GFP (F) or Vinc::GFP (G) and E-Cad at st. 13–16. As controls, M2 coefficient is also calculated using clockwise-rotated GFP signals at st. 14. A two-tailed non-parametric Mann–Whitney test between st.14 and 90°-rotated st.14 showed no significant difference (ns). Scale bars for all panels 10 μm.

### Fig. S2 Carvalho et al.

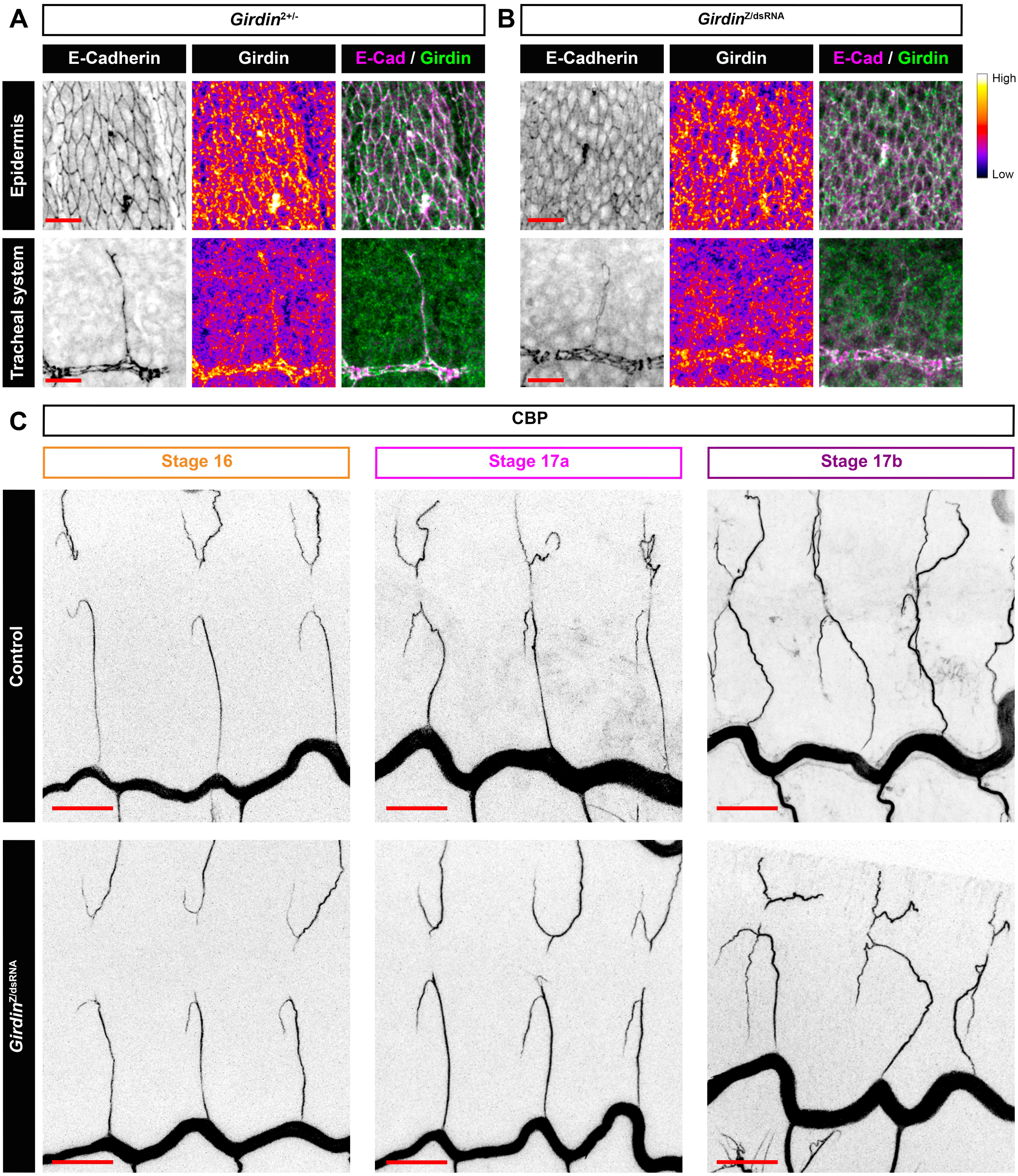


**Figure S2. Girdin distribution under Girdin-depleted conditions and interspace between fusion cells in control and *Girdin* $^{Z/dsRNA}$ embryos.**

(A,B) Girdin distribution at AJs in epidermal and DB tracheal cells in *Girdin*$^{2+/-}$ heterozygote mutants (A) and *Girdin*$^{Z/dsRNA}$ embryos (B) at st. 14. E-Cad and Girdin signals are shown as inverted and Fire LUTs, respectively. Scale bars: 10 µm. (C) Interspace between fusion cells revealed by CBP staining (shown as inverted LUT) at st. 16, 17a and 17b in control and *Girdin*$^{Z/dsRNA}$ tracheal cells. St. 17 is divided into two parts based on the terminal lumen length: st. 17a (10-20 µm) and st. 17b (>20 µm). Scale bars: 20 µm.

### Fig. S3 Carvalho et al.

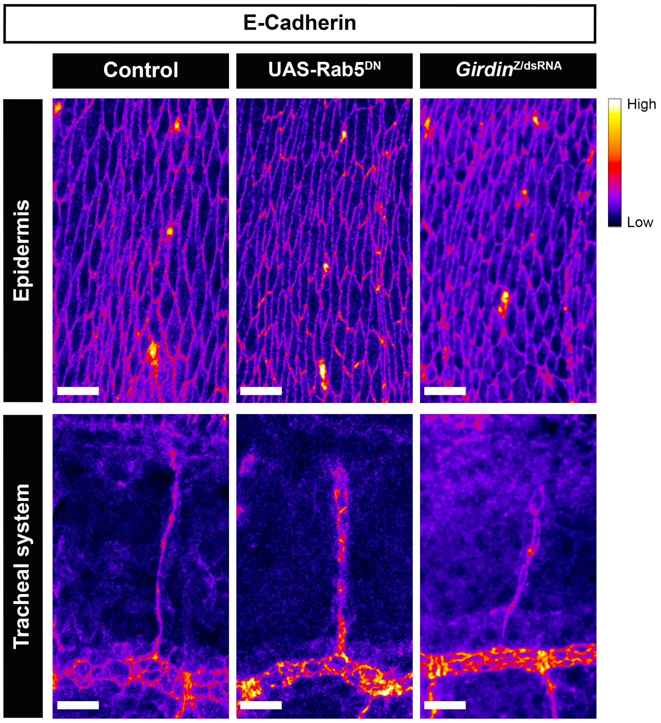


**Figure S3. E-Cad distribution in controls and *Girdin* $^{Z/dsRNA}$ mutants.**

E-Cad levels shown as Fire LUTs, in the epithelium and the corresponding DB under control, UAS-Rab5$^{DN}$ and *Girdin*$^{Z/dsRNA}$ conditions. Labelling in epithelial cells serves as an internal reference, since UAS-*Girdin*-dsRNA is not expressed in this tissue. Scale bars, 10 µm.

### Fig. S4 Carvalho et al.

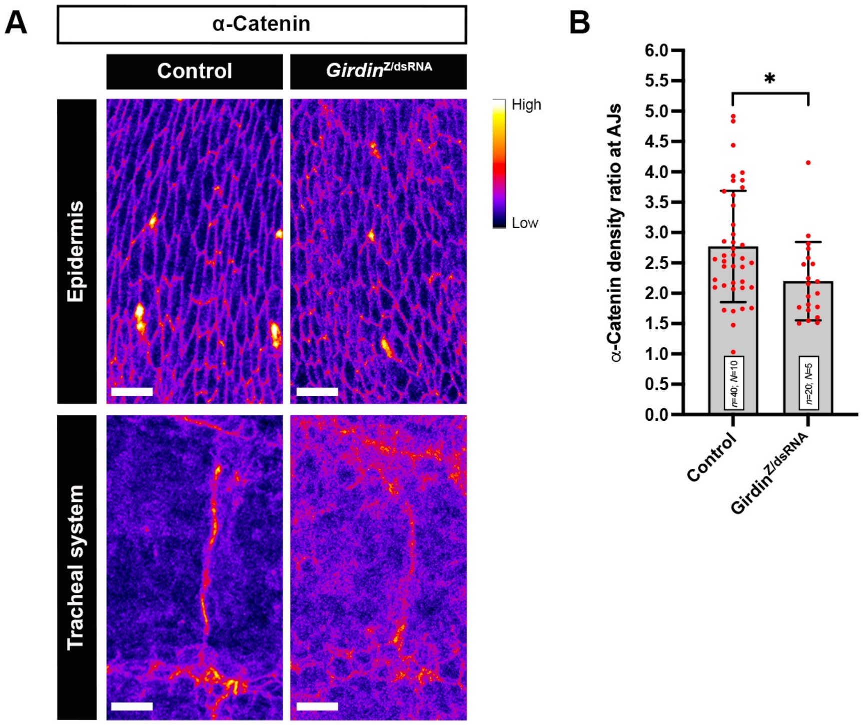


**Figure S4. α-Catenin density ratios at AJs in control and *Girdin*$^{Z/dsRNA}$ DBs compared to epidermal cells.**

(A) α-Catenin levels shown as Fire LUTs, in the epidermis and the corresponding DB under control and *Girdin*$^{Z/dsRNA}$ conditions. Labelling in epidermal cells serves as an internal reference, since UAS-*Girdin*-dsRNA is not expressed in this tissue. Scale bars, 10 µm. (B) Ratios were calculated between α-Catenin signal densities at AJs of DBs and those of epidermal cells in control and *Girdin*$^{Z/dsRNA}$ embryos. A two-tailed non-parametric Mann–Whitney test between controls and *Girdin*$^{Z/dsRNA}$ DBs shows significant difference (*, $p < 0.05$).

Fig. S5 Carvalho et al.

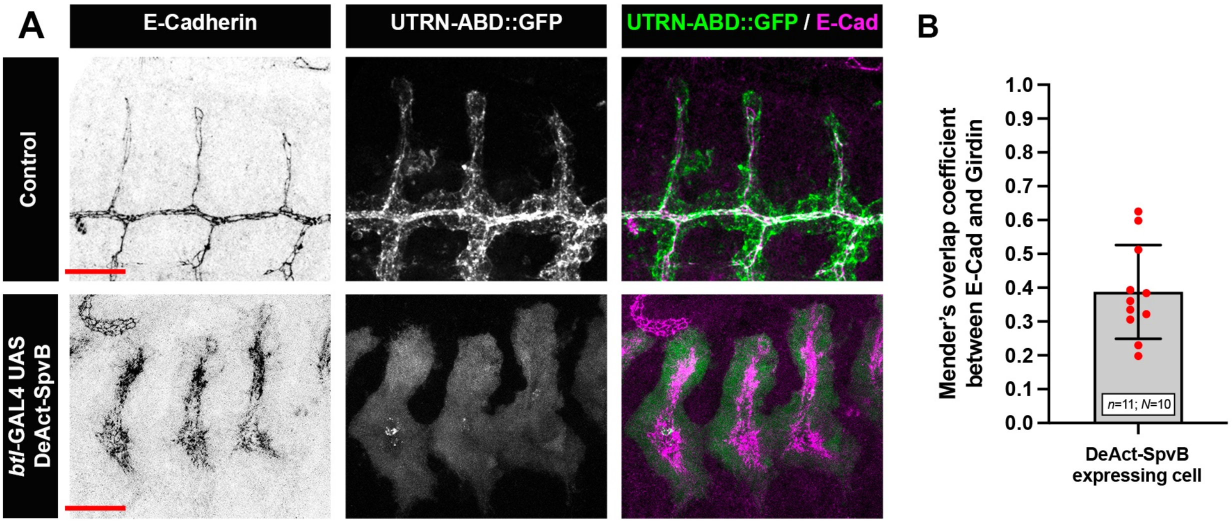


**Figure S5. Effects of DeAct-Spvb expression on GFP–UTRN::ABD actin probe distribution and Girdin localisation to AJs.**

(A) GFP–UTRN::ABD expression in control and DeAct-SpvB–expressing tracheal cells. AJs are marked by E-Cad. The diffuse GFP signal represents cytoplasmic GFP–UTRN::ABD unbound to F-actin. Scale bars, 20 µm. (B) Manders' overlap coefficient (M2) between Girdin and E-Cad at st. 14 in DeAct-SpvB–expressing tracheal cells.
